# A 3D *in vitro* follicle-endometrial organoid co-culture model recapitulates ovarian control of human endometrial decidualization

**DOI:** 10.64898/2026.09.24.754184

**Authors:** Yubing Liu, Yang Sun, Tingjie Zhan, Xueyuan Liu, Jiyang Zhang, Feng Gao, Nataki C. Douglas, Qiang Zhang, Shuo Xiao

## Abstract

Female infertility remains a major women’s reproductive health challenge, with impaired endometrial receptivity contributing to recurrent implantation failure, early pregnancy loss, and pregnancy complications. Mechanistic studies of human early pregnancy are limited by the lack of physiologically relevant models, and existing *in vitro* endometrial models rely on exogenous hormone treatment and do not recapitulate the dynamic ovarian control of endometrial growth and differentiation. Here, as a proof of concept, we developed a 3D *in vitro* ovarian follicle-endometrial organoid co-culture model. Immature murine follicles were cultured *in vitro* to obtain preovulatory follicles, followed by human chorionic gonadotropin (hCG) stimulation to induce ovulation and luteinization. Luteinized follicles were co-cultured with human endometrial stroma organoids for 6 days. The results showed that luteinized follicles generated a dynamic endocrine environment characterized by progressively increasing progesterone levels and declining estradiol levels. Follicle co-culture induced morphological differentiation and expression of decidualization markers in human endo-organoids. Transcriptomic analysis revealed strong genome-wide concordance between follicle co-culture and conventional decidualization induction using estradiol/progesterone/cAMP (EPC), with significant enrichment of decidualization-associated genes and pathways under both conditions. Nevertheless, the two conditions also exhibited distinct transcriptional states, with EPC preferentially enriched for more mature decidualization-associated programs, whereas luteinized follicle co-culture retained features of the early stage of decidualization. Functional perturbation testing revealed that the progesterone receptor (PR) antagonist RU486 attenuated co-culture-induced endometrial decidualization. Together, these results establish a new *in vitro* new approach methodology (NAM) that recapitulates ovarian control of endometrial decidualization. This platform provides a human-relevant NAM model to investigate the physiology and pathophysiology of endometrial decidualization, and to evaluate how drugs and environmental chemicals perturb uterine receptivity and early pregnancy.

## Introduction

The ovary and uterus function as an integrated reproductive axis in which temporally coordinated ovarian signals regulate the cyclic remodeling of the endometrium and establish an environment conducive to embryo implantation, placentation, and pregnancy. During the follicular phase, increasing estradiol (E2) production by developing ovarian follicles promotes endometrial growth. Following ovulation, luteinized follicles give rise to the corpus luteum (CL), which primarily produces progesterone (P4) and other CL-derived paracrine factors. These signals transform the estrogen-primed endometrium into a receptive, secretory tissue and promote decidualization, thereby supporting embryo implantation and other early pregnancy events before placental formation [1].

Dynamic changes in E2 and P4 orchestrate the progressive remodeling of endometrial stromal cells into secretory decidual cells. This profound decidualization process is characterized not only by the induction of canonical markers and key regulators, such as Insulin-like growth factor-binding protein 1 (*IGFBP1)*, prolactin (*PRL),* and Forkhead box protein O1 (*FOXO1*), but also by extensive extracellular matrix (ECM) remodeling, inflammatory-like responses, and metabolic reprogramming. Beyond steroid hormones, the CL produces a diverse repertoire of steroid metabolites, prostaglandins, peptides, and vasoactive and angiogenic factors that directly or indirectly influence the endometrium. These findings suggest that the physiological regulation of the endometrium by the ovary involves a complex and dynamically evolving secretory environment [2].

*In vitro* decidualization can be induced in primary or immortalized human endometrial stromal cells (HESCs) by treatment with defined concentrations of E2, P4, and/or cAMP, commonly termed EPC, providing fundamental insights into the cellular and molecular mechanisms underlying this process [3–6]. Emerging three-dimensional (3D) endometrial culture and organoid models provide a more physiologically relevant extracellular environment and tissue architecture for studying decidualization [7–10]. We recently demonstrated that hormonal stimulation with EPC induces a broader decidualization-associated transcriptional response in 3D human endometrial organoids than cAMP alone [11]. Nevertheless, these approaches depend on exogenous hormones and signaling agonists at predefined concentrations. Although highly useful for mechanistic studies, they do not recapitulate the temporally changing endocrine output of a functional ovarian compartment, such as CL, or directly investigate how ovarian follicle development, ovulation, and luteinization are coordinated with endometrial growth and differentiation. Thus, a major gap in current *in vitro* female reproductive models is the lack of a functionally coupled ovary–endometrium system that recapitulates physiological inter-organ communication.

To address this knowledge gap, we developed a 3D *in vitro* follicle–endometrial organoid co-culture model integrating our established *in vitro* follicle growth (IVFG) system with human endometrial organoids. Our previous studies have demonstrated that the IVFG supports follicular growth, ovulation, and luteinization, recapitulating key ovarian endocrine functions [12–27]. Preovulatory follicles grown from IVFG were stimulated with human chorionic gonadotropin (hCG) to induce ovulation and luteinization and then co-cultured with human endometrial organoids, allowing them to respond to endogenous and temporally changing ovarian signals. We hypothesized that this functional ovarian compartment would induce human endometrial decidualization with molecular responses overlapping with, but also distinct from, the conventional EPC stimulation. We characterized ovarian hormone production, endometrial morphological and molecular responses, and transcriptomic profiles, and performed pharmacological perturbations to evaluate the functional utility of the co-culture model. Together, this study establishes a multi-organ 3D *in vitro* culture model to investigate ovarian control of endometrial decidualization and the effects of pharmacological and environmental chemicals on ovary–endometrium communication, reproductive cycles, and early pregnancy.

## Materials and Methods

### Animals

CD-1 mice used in this study were obtained from our in-house breeding colony established from animals purchased from Envigo (Envigo, Indianapolis, IN) and maintained at Rutgers University. Mice were housed under controlled conditions (22 ± 1°C, 30%–70% humidity) on a 12-h light/12-h dark cycle. All animal procedures were conducted in accordance with the National Institutes of Health Guide for the Care and Use of Laboratory Animals and were approved by the Rutgers University Institutional Animal Care and Use Committee (IACUC).

### 3D *in vitro* follicle growth (IVFG)

Immature preantral follicles were isolated from the ovaries of 16-day-old female CD-1 mice and cultured using our established 3D *in vitro* follicle growth (IVFG) system. Briefly, ovaries were collected following CO₂ euthanasia and incubated in L-15 medium (Invitrogen, Carlsbad, CA) containing Liberase (Sigma-Aldrich, St. Louis, MO) and DNase I to facilitate follicle isolation. Immature preantral follicles at the secondary stage, measuring approximately 150–180 μm in diameter, were manually isolated under a stereomicroscope and individually encapsulated in 0.5% alginate hydrogel (Sigma-Aldrich, St. Louis, MO). Encapsulated follicles were cultured individually in 96-well plates in follicle growth medium supplemented with 10 mIU/mL recombinant FSH (rFSH; Organon; provided by Dr. Mary Zelinski, Oregon National Primate Research Center, Oregon Health & Science University, Beaverton, OR, USA). Follicles were maintained in culture until they reached the preovulatory stage.

### Human endometrial stromal cell (HESC) *in vitro* culture and 3D endometrial organoid generation

Primary human endometrial stromal cells previously established and characterized by the Douglas laboratory at Rutgers University were used in this study. The original human endometrial tissues were obtained from consented donors under Rutgers University IRB protocol Pro2018002041[11]. As shown in Figure 1A, HESCs were cultured at 37°C in a humidified atmosphere containing 5% CO_₂_in phenol red-free Dulbecco’s modified Eagle’s medium/Ham’s F-12 (DMEM/F12; 1:1) supplemented with 10% charcoal/dextran-treated fetal bovine serum, 1% ITS+ Premix, 500 ng/mL puromycin, and 1% penicillin–streptomycin. For 3D endometrial organoid formation, agarose microwell hydrogels were fabricated using reusable 3D Petri Dish® 24-96 Small Spheroid silicone micro-molds (Microtissues, Inc.; SKU 24-96). Molten agarose was cast into the silicone micro-molds and allowed to solidify. The resulting agarose hydrogels containing 96 round-bottomed microwells were removed from the molds and placed into 24-well culture plates, with one microwell per well. After equilibration with culture medium, HESCs were dissociated into a single-cell suspension, and approximately 2 × 10 cells were seeded onto each agarose hydrogel insert. Cells were cultured in MammoCult spheroid medium for 24 h to allow self-assembly into organoids, with approximately 80-90 organoids in each agarosemicrowell. The MammoCult medium was then completely replaced with HESC maintenance medium for subsequent experiments.

**Figure 1.**
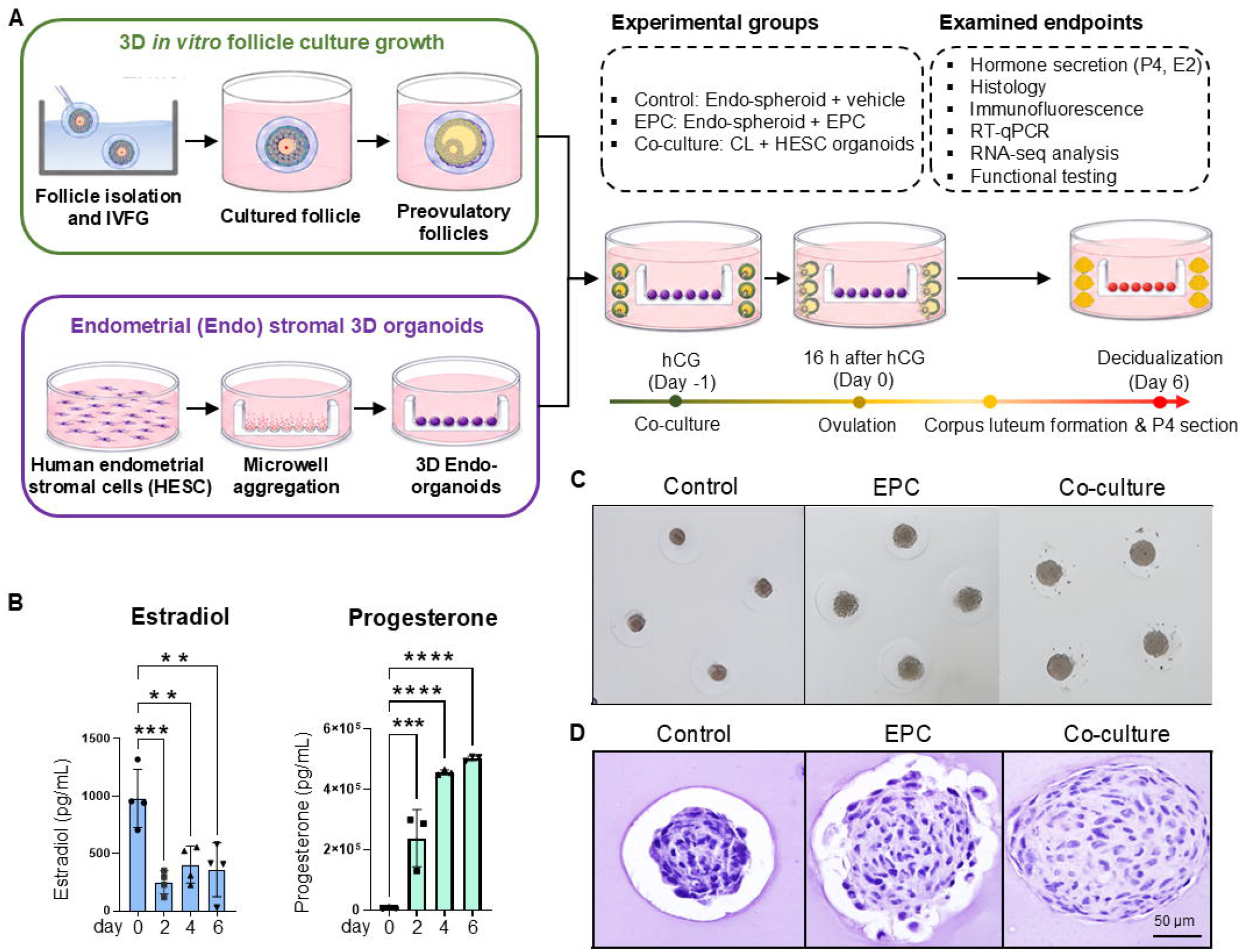
Establishment of the follicle–endometrial organoid co-culture model and induction of endometrial decidualization. **(A)** Schematic of the 3D *in vitro* follicle–endometrial organoid co-culture model. Immature mouse ovarian follicles were isolated and cultured using the *in vitro* follicle growth (IVFG) system to generate preovulatory follicles. Human endometrial stromal cells (HESCs) were aggregated in agarose microwells to generate 3D endometrial organoids. Preovulatory follicles were co-cultured with endometrial organoids and stimulated with human chorionic gonadotropin (hCG) to induce ovulation and luteinization. Sixteen hours after hCG stimulation was designated day 0, and co-culture was continued through day 6. Endometrial organoids cultured with vehicle or estradiol/progesterone/cAMP (EPC) served as negative and positive controls, respectively. **(B)** Estradiol (E2) and progesterone (P4) concentrations in conditioned medium during follicle co-culture. **(C)** Representative bright-field images of endometrial organoids after 6 days of vehicle, EPC, or follicle co-culture treatment. Scale bar, 200 μm. **(D)** Representative hematoxylin and eosin (H&E)-stained sections of endometrial organoids after 6 days of treatment. Scale bar, 50 μm. Data are presented as mean ± SD. Statistical significance was determined by one-way repeated-measures ANOVA followed by Dunnett’s multiple-comparisons test using day 0 as the reference. \*\**P < 0.01*, \*\*\**P < 0.001*, \*\*\*\**P < 0.0001*.

### Decidualization induction and follicle–endometrial organoid co-culture

Following 24 h of organoid formation in MammoCult medium, the medium was replaced with HESC culture medium. Preovulatory follicles grown from IVFG were then combined with endometrial organoids. Six preovulatory follicles were added to each agarose microwell containing 96 endometrial organoidsand stimulated with 1.5 IU/mL hCG for 16 h to induce ovulation and luteinization. Following hCG stimulation, the ovulation medium was replaced with endometrial organoid maintenancemedium, and this time point was designated as co-culture day 0. Endometrial organoids were maintained for 6 days under three experimental conditions: (1) vehicle control; (2) EPC consisting of 10 nM E2, 1 μM P4, and 0.5 mM cAMP as a positive control for decidualization induction; or (3) follicle co-culture, in which 80-90 endometrial organoids were co-cultured with six luteinized follicles per well. Conditioned medium was collected at the indicated time points for hormone and secreted-factor measurements, and endometrial organoids were collected after 6 days for histological, molecular, and transcriptomic analyses.

### Measurement of P4, E2, and PGE2

E2, and P4 concentrations in conditioned culture medium were quantified using commercially available ELISA kits from Cayman Chemical (Ann Arbor, MI, USA) according to the manufacturer’s protocols. P4 was measured using the kit of Cat. No. 582601 and E2 using Cat. No. 501890. Conditioned medium was stored at −20°C until analysis. For each assay, conditioned medium and serially diluted standards were added to the assay plate together with the corresponding antibody and acetylcholinesterase (AChE)-linked tracer. Following the recommended incubation period, plates were washed to remove unbound components, and Ellman’s reagent was added for color development. Optical density was measured using a SpectraMax M3 microplate reader (BioTek Instruments, Winooski, VT, USA) at 414 nm for P4 and E2. Hormone concentrations were calculated from the corresponding standard curves. The assay ranges were 7.8–1,000 pg/mL for P4 and 0.61–10,000 pg/mL for E2. Conditioned-media samples were diluted when necessary to ensure that measured values fell within the linear range of the standard curve. Standards and assay blanks were included in each experiment for quality control.

### Histology and immunofluorescence staining

At the end of 6 days of co-culture, endometrial organoids were collected and fixed overnight in 4% paraformaldehyde (PFA). Fixed organoids were processed for paraffin embedding and sectioned at a thickness of 5 μm. For histological evaluation, sections were deparaffinized, rehydrated, and stained with hematoxylin and eosin (H&E) using standard procedures.

For immunofluorescence staining, paraffin sections were deparaffinized and rehydrated, followed by antigen retrieval and blocking. Sections were incubated overnight at 4°C with a rabbit polyclonal anti-IGFBP1 primary antibody (Abcam, Cambridge, MA, USA; Cat. No. ab228741; [1:1000]). After washing, sections were incubated with a Goat anti-Rabbit IgG (H+L) Highly Cross-Adsorbed Secondary Antibody, Alexa Fluor™ Plus 488 (Thermo Fisher Scientific, Waltham, MA, USA; Cat. No. A32731TR; 1:5000). Slides were mounted with VECTASHIELD antifade mounting medium containing DAPI (Maravai LifeSciences, San Diego, CA, USA) and imaged using a Leica confocal microscope (Leica Microsystems, Wetzlar, Germany).

### RNA extraction and RT-qPCR

Endometrial organoids from each culture well were collected on day 6 and pooled for total RNA extraction using the Arcturus PicoPure RNA Isolation Kit (Applied Biosystems) according to the manufacturer’s instructions, followed with RT-qPCR as we described [25]. Supplementary Table 1 provides the names, functions, and relevant references for these genes, while the corresponding primer sequences are provided in Supplementary Table 2.

### RNA sequencing of endometrial organoids and bioinformatic analysis

The endometrial organoid total RNA was used for low-input RNA library preparation and sequencing by Novogene Corporation (Sacramento, CA, USA) on an Illumina NovaSeq platform using 150-bp paired-end sequencing.

Fifteen biological samples were included, consisting of four vehicle control samples, five EPC-treated samples, and six follicle co-culture samples. Sequencing reads were processed in Partek Flow. Adapter sequences and low-quality bases were removed before alignment, and reads corresponding to ribosomal and mitochondrial sequences were filtered using Bowtie 2. The remaining reads were aligned to the human reference genome GRCh38. Gene-level read counts were generated using Ensembl transcript annotations (release 99) and the Partek expectation-maximization algorithm. Transcript abundance was additionally expressed as transcripts per million (TPM), and analyses were restricted to protein-coding genes based on HUGO Gene Nomenclature Committee annotations.

Principal component analysis (PCA) was performed using DESeq2 and plotted in R with ggplot2 to visualize global transcriptional relationships among vehicle control, EPC, and co-culture samples. Differential gene expression was analyzed using DESeq2 from raw gene counts. To evaluate the overall similarity of transcriptional responses induced by EPC treatment and follicle co-culture, gene-wise log_2_ fold changes from the EPC-versus-control and co-culture-versus-control comparisons were directly compared. Pearson correlation analysis was used to quantify concordance between the two transcriptional responses. Genes were further categorized according to statistical significance and direction of regulation in the two comparisons as shared, condition-specific, oppositely regulated, or non-significantly regulated.

Gene set enrichment analysis (GSEA) was performed using clusterProfiler in R to assess coordinated decidualization-associated transcriptional responses. A previously defined “*In vitro* decidualization markers” gene set, containing established decidualization markers and additional genes associated with in vitro decidualization, was evaluated separately in the EPC-versus-control and co-culture-versus-control comparisons. Additional decidualization-related pathways were grouped into functional categories representing endocrine and vascular signaling, immune and inflammatory responses, lipid and energy metabolism, and cell–cell and cell–matrix interactions. Enrichment was summarized using normalized enrichment scores (NES) and corresponding significance values.

To compare the relative decidualization states of the EPC and co-culture models, the previously defined *in vitro* decidualization marker set was further stratified according to early-secretory, mid- to late-secretory, and first-trimester-associated markers. Enrichment of these stage-associated gene sets was evaluated in the EPC-versus-co-culture comparison. In addition, transcriptional staging was performed using the Endest model as a reference-based estimate of relative endometrial transcriptional state. Because Endest was originally developed using human endometrial tissue, the resulting scores were interpreted comparatively rather than as absolute histological staging.

Finally, pathway enrichment analysis was performed on transcriptional differences between the EPC and co-culture groups to identify biological programs preferentially associated with each model. Pathways related to decidual maturation, hormone signaling, metabolic regulation, inflammatory and growth-factor signaling, and cellular proliferation were examined to characterize shared and condition-specific transcriptional features.

### RU486 treatment and functional testing

To demonstrate the utility of the co-culture model for functional testing, endometrial organoids were treated with the PR antagonist mifepristone (RU486; 10 μM) during the 6-day decidualization induction period. RU486 was added to both the follicle co-culture and EPC groups, with corresponding vehicle-treated groups included as controls. At the end of treatment, endometrial organoids were collected for morphological evaluation and RT-qPCR analysis of decidualization-associated genes.

### Statistical analysis

Statistical analyses were performed using GraphPad Prism [10.1.0 (GraphPad Software, Boston, MA, USA). Data are presented as mean ± standard deviation (SD) unless otherwise indicated. Biological replicates were considered independent experimental units. For longitudinal hormone measurements obtained from the same cultures at multiple time points, differences over time were analyzed using one-way repeated-measures analysis of variance (ANOVA), followed by Dunnett’s multiple-comparisons test using the initial time point as the reference. For comparisons among the vehicle control, EPC, and co-culture groups, one-way ANOVA followed by Dunnett’s multiple-comparisons test was used as appropriate. A two-sided *P* value < 0.05 was considered statistically significant. Statistical procedures specific to the RNA-seq analyses, including differential gene expression and gene set enrichment analyses, are described in the RNA-seq and bioinformatic analysis section. False discovery rate (FDR)-adjusted *P* values were used where applicable.

## Results

### Follicle co-culture recapitulates luteal endocrine dynamics and promotes endometrial decidualization

We established an *in vitro* follicle–endometrial organoid co-culture model by combining preovulatory follicles generated using our IVFG system with human endometrial organoids (Figure 1A). Preovulatory follicles were stimulated with hCG to induce ovulation and luteinization and subsequently maintained in co-culture with endometrial organoids at day 0. Endometrial organoids cultured alone served as controls, whereas endometrial organoids treated with EPC served as the positive control. Following hCG stimulation, E2 and P4 levels in the coculture medium showed dynamic changes that closely resembled those observed *in vivo*. Estradiol peaked before ovulation, which occurred 16 h after hCG stimulation, then rapidly decreased before rising again to a moderate level. Progesterone gradually increased following ovulation (Figure 1B).

Bright-field imaging showed that during the 6-day co-culture period, endometrial organoids treated with vehicle remained relatively small, compact, and spherical, whereas endometrial organoids treated with EPC or co-cultured with luteinized follicles appeared larger and exhibited a less compact morphology (Figure 1C). These differences were more evident by H&E staining (Figure 1D). Control endometrial organoids consisted of densely packed cells with limited cytoplasm. In contrast, endometrial organoids treated with EPC or co-cultured with luteinized follicles showed a significant expansion in organoid size, along with more cytoplasmic and extracellular matrix structures. These morphological features are consistent with the transformation of fibroblast-like endometrial stromal cells toward an enlarged decidual cell phenotype. Together, these results suggest that the follicle–endometrial organoid co-culture system generates dynamic luteal endocrine signals and promotes morphological differentiation characteristic of endometrial decidualization.

### Follicle co-culture induces molecular features of endometrial decidualization

To determine whether the morphological changes induced by follicle co-culture were accompanied by molecular features of endometrial decidualization, we next examined the expression of decidualization marker genes in endometrial organoids. As expected, the results of RT-qPCR showed that EPC significantly increased the expression of decidualization markers of *IGFBP1, PRL, LEFTY, FOXO1, DKK1, DCN, REN,* and *SST* compared with untreated controls (Figure 2A). Although the magnitude of transcriptional induction varied among individual genes, endometrial organoids co-cultured with luteinized follicles exhibited similar transcription induction of these genes (Figure 2A). Consistent with the RT-qPCR results, immunofluorescence staining showed minimal IGFBP1 expression in untreated endometrial organoids, whereas EPC and follicle co-culture significantly increased IGFBP1 expression (Figure 2B). Together with the morphological changes, these findings demonstrate that luteinized follicles induce molecular features of human endometrial organoid decidualization.

**Figure 2.**
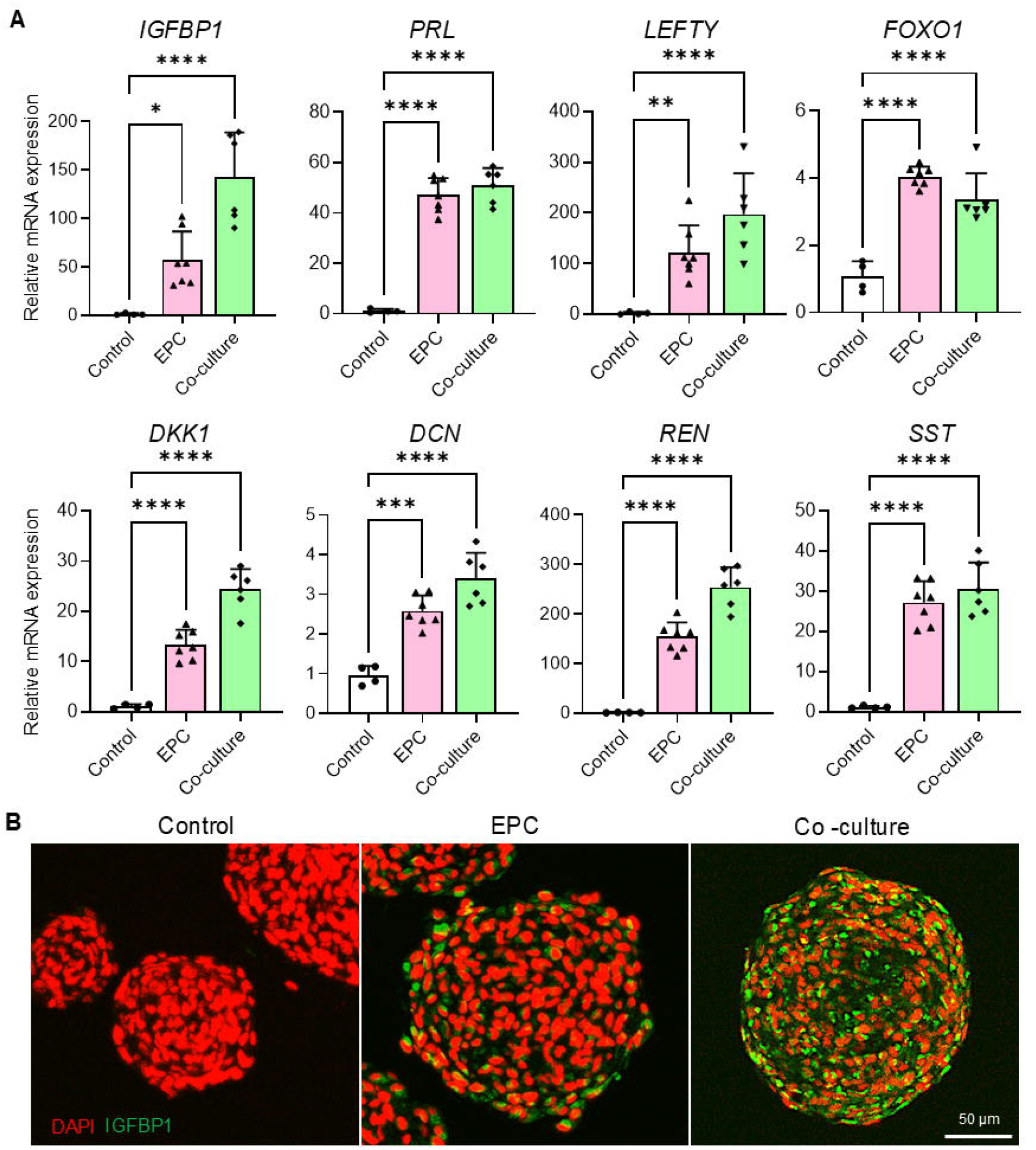
Follicle co-culture induces molecular features of endometrial decidualization. **(A)** RT-qPCR analysis of decidualization-associated genes, including *IGFBP1, PRL, LEFTY, FOXO1, DKK1, DCN, REN,* and *SST*, in human endometrial organoids following 6 days of vehicle control, EPC treatment, or co-culture with luteinized follicles. Gene expression was normalized to the vehicle control group. **(B)** Representative immunofluorescence images showing IGFBP1 expression in endometrial organoids following vehicle, EPC, or follicle co-culture treatment. IGFBP1 is shown in green, and nuclei are counterstained with DAPI (red). Scale bar, 50 μm. Data are presented as mean ± SD. Statistical significance was determined by one-way ANOVA followed by Dunnett’s multiple-comparisons test. \**P < 0.05*, \*\**P < 0.01*, \*\*\**P < 0.001*, \*\*\*\**P < 0.0001*.

### Follicle co-culture induces a comparable transcriptional profile with EPC treatment

RNA-seq analysis was performed to examine transcriptomic changes in endometrial organoids in an unbiased manner. PCA revealed clear separation of organoids treated with EPC or co-cultured with luteinized follicles from organoids treated with vehicle, whereas the difference between the co-culture and EPC groups was relatively smaller (Figure 3A). Sample similarity analysis showed within-group consistency across all groups, with the EPC and co-culture samples displaying relatively small intergroup distances and clustering closely together (Figure 3B).

**Figure 3.**
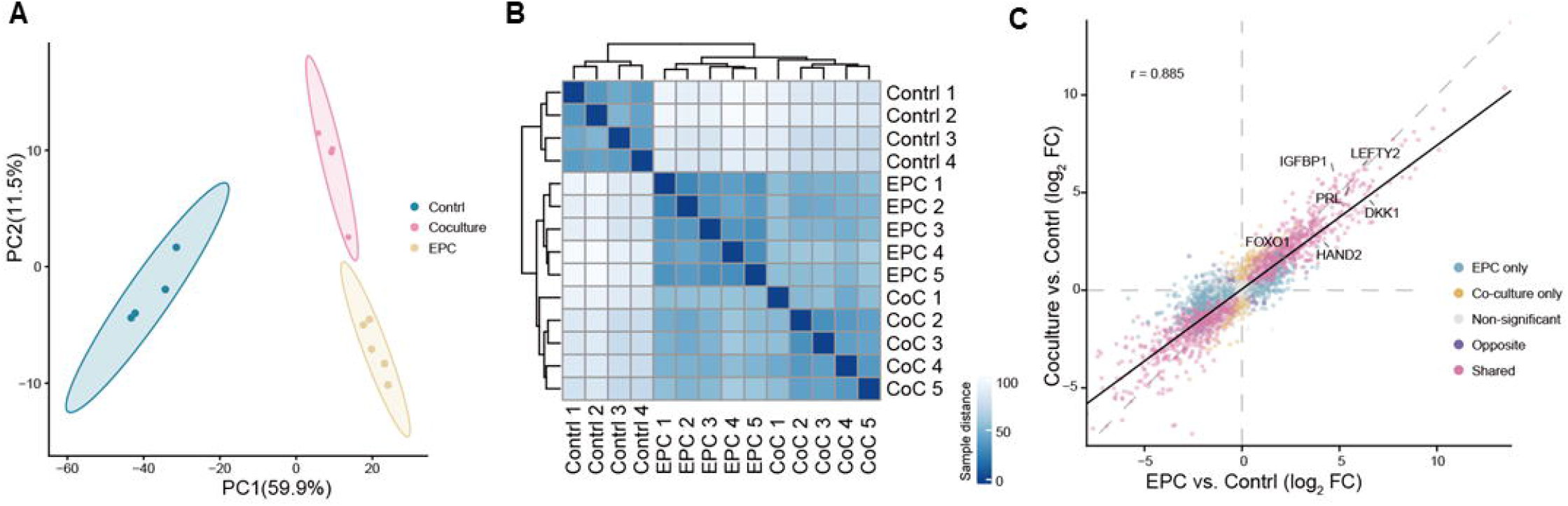
Follicle co-culture induces a global transcriptional response comparable to conventional EPC-induced decidualization. RNA-seq was performed on human endometrial organoids treated with vehicle control (n = 4), EPC (n = 5), or follicle co-culture (n = 6). **(A)** Principal component analysis (PCA) showing global transcriptional relationships among control, EPC, and follicle co-culture groups. **(B)** Sample-to-sample similarity analysis illustrating within- and between-group transcriptional relationships. **(C)** Comparison of gene-wise log2 fold changes (FC) for EPC versus control and follicle co-culture versus control. Each point represents an expressed gene and is categorized according to whether it was significantly altered by EPC only, co-culture only, both conditions (shared), oppositely regulated, or not significantly altered. Selected decidualization-associated genes are indicated. The Pearson correlation coefficient (*r* = 0.885) indicates strong concordance between transcriptional responses induced by EPC and follicle co-culture.

To further evaluate the overall similarity of transcriptomic responses induced by EPC and follicle co-culture, we compared gene-wise log2 fold changes relative to the control group. Across expressed genes, the transcriptomic responses showed a markedly high degree of concordance, with a Pearson correlation coefficient of 0.885 (Figure 3C). Most significantly altered genes, including highlighted decidualization markers of IGFBP1, PRL, LEFTY2, FOXO1, HAND2, and DKK1, exhibited consistent directions and broadly similar magnitudes of transcriptional change between EPC and co-culture groups, with only a small subset of genes showing discordance (Figure 3C).

### Follicle co-culture recapitulates decidualization-related transcriptomic changes in endometrial organoids

We next performed GSEA using a gene set termed “*In vitro* decidualization markers” for the EPC vs. Control and Co-culture vs. Control comparisons. This gene set was previously defined based on established decidualization markers and additional genes associated with *in vitro* decidualization [28]. The results revealed that these markers were significantly enriched in both comparisons, with comparable enrichment levels (Figure 4A).

**Figure 4.**
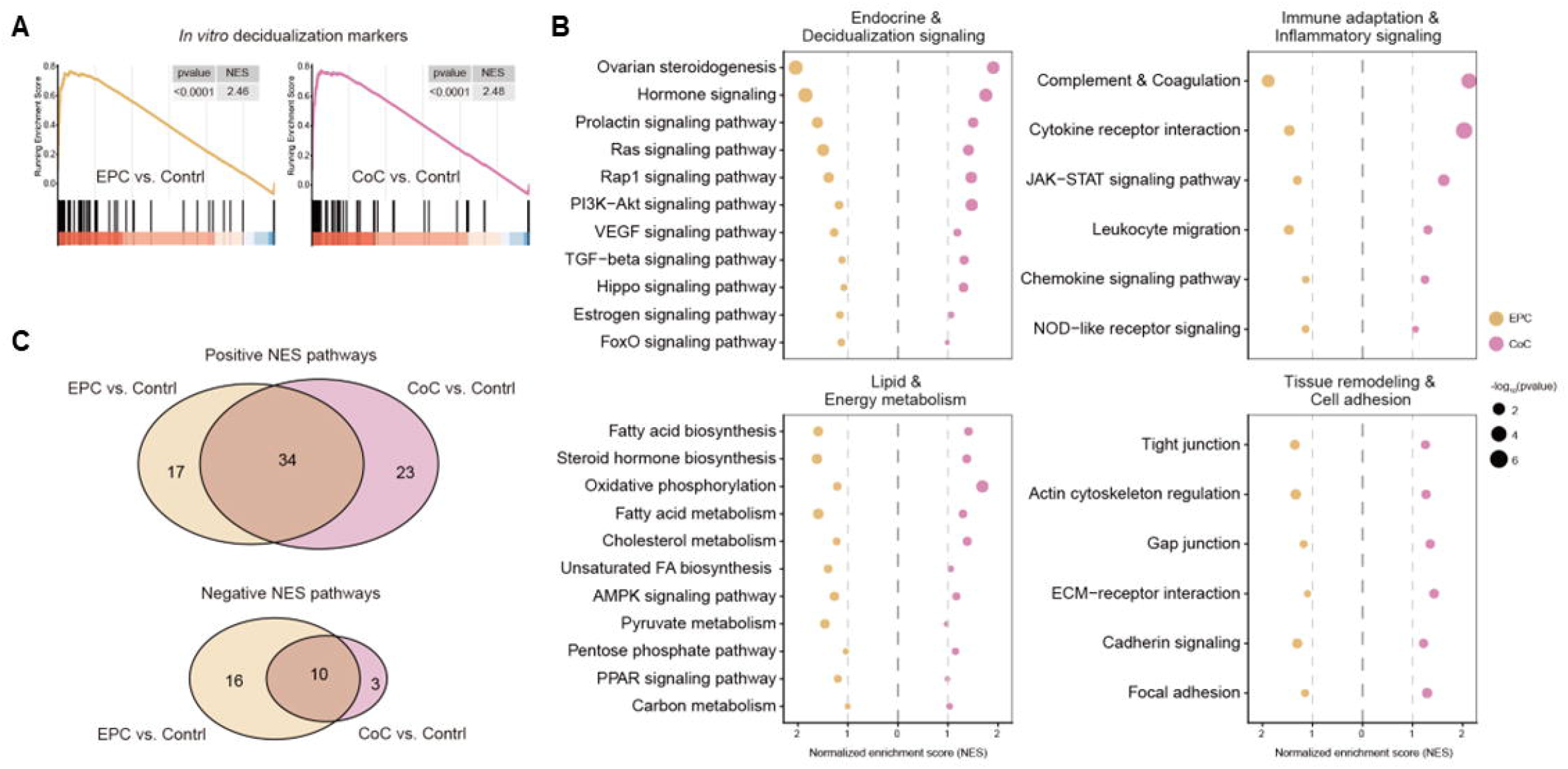
Follicle co-culture recapitulates decidualization-associated transcriptional programs induced by EPC. **(A)** Gene set enrichment analysis (GSEA) of the previously defined “*In vitro* decidualization markers” gene set in EPC versus control and follicle co-culture versus control comparisons. Normalized enrichment scores (NES) and nominal *P* values are shown. **(B)** Comparative pathway enrichment analysis of decidualization-associated biological programs grouped into four functional categories: endocrine and decidualization signaling, immune adaptation and inflammatory signaling, lipid and energy metabolism, and tissue remodeling and cell adhesion. Dot position represents NES, and dot size represents statistical significance as −log10(*P* value). **(C)** Venn diagrams showing the numbers of significantly positively and negatively enriched pathways shared between or unique to EPC versus control and follicle co-culture versus control comparisons.

As decidualization is a coordinated tissue-remodeling process encompassing multiple physiological programs, we further examined decidualization-related cellular and molecular programs across four functional categories in both comparisons (Figure 4B, Supplemental Table 4). Endocrine and decidualization-related signaling pathways were consistently enriched in both comparisons, encompassing hormonal responsiveness, vascular signaling, and cAMP-associated regulation. Immune adaptation and inflammatory signaling were also consistently enriched in both comparisons, encompassing cytokine-, chemokine-, and innate immune-related programs characteristic of the inflammatory-like microenvironment of the decidua. Metabolic programs associated with decidualization were similarly enriched, encompassing coordinated changes in lipid and energy metabolism. Lastly, pathways involved in cell–cell and cell–matrix interactions were also enriched, consistent with tissue remodeling accompanying the decidualization process. Beyond decidualization-related programs, both EPC and follicle co-culture showed substantially more positive than negative enrichment relative to the control (Figure 4C), indicating a broad induction of transcriptional programs under both conditions. Nevertheless, each condition also displayed a subset of uniquely enriched pathways, indicating that follicle co-culture and EPC induce broadly similar but not identical transcriptional responses. Collectively, these transcriptomic analyses indicate that follicle co-culture recapitulates major decidualization transcriptional responses induced by conventional EPC treatment while retaining distinct molecular features.

### The follicle co-culture model induces early stage of endometrial decidualization than the EPC

To identify the major transcriptional differences between the EPC and follicle co-culture groups, we performed differential expression analysis and examined the expression of *in vitro* decidualization markers. According to the original report, this gene set comprises markers associated with the early-, mid- to late-secretory, and first-trimester stages of decidualization [28]. Among genes more highly expressed in EPC, several markers associated with first-trimester decidua, including *SLC1A1*, *HSD11B1*, *SCARA5*, *FOXO1*, *HAND2*, *APOD*, and *PROK1*, as well as late-secretory markers such as *OSR2*, *P4HA2*, and *DDIT4* showed significant differences between the two conditions (Figure 5A, Supplemental Table 5). In contrast, *VEGFA*, *SEMA3A*, *MMP11*, and *WNT5A*, which are associated with early secretory-phase endometrium, were highly expressed in endometrial organoids treated with luteinized follicles (Figure 5A, Supplemental Table 5). Consistent with these differences, mid- to late-secretory and first-trimester decidualization markers were preferentially enriched in EPC, with significant enrichment for first-trimester markers (Figure 5B). These findings are consistent with conventional EPC-induced decidualization preferentially acquiring features of more mature, pregnancy-associated decidua [28]. However, the original marker set was biased toward later stages, particularly the first-trimester decidualization, and contained relatively few early-stage markers. Thus, the lack of significant enrichment of the early-stage gene set does not exclude early decidualization features in the co-culture group.

**Figure 5.**
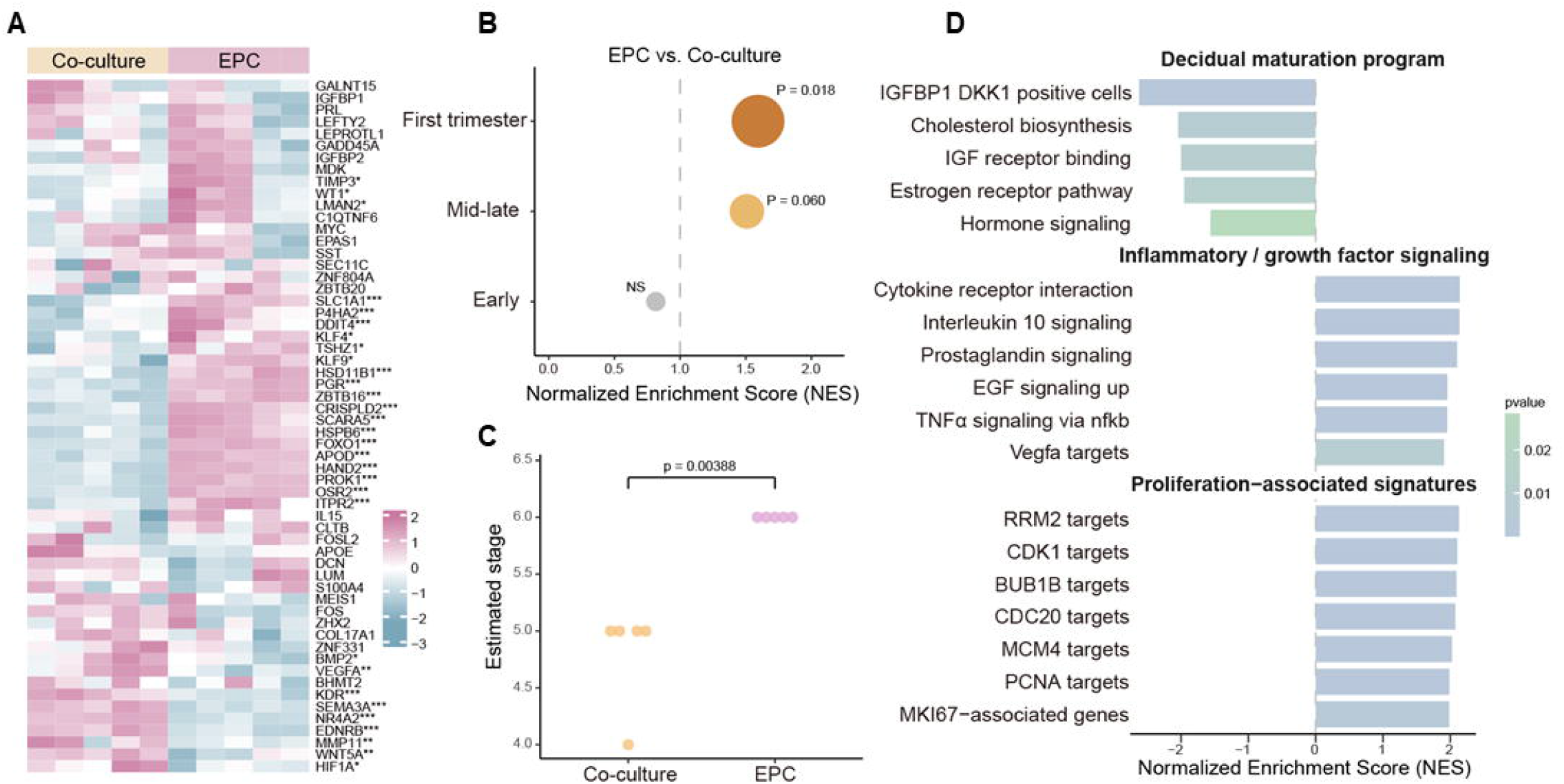
Follicle co-culture and EPC induce distinct decidualization-associated transcriptional states. **(A)** Heatmap showing expression of previously defined *in vitro* decidualization markers associated with first-trimester, mid- to late-secretory, and early-secretory endometrial states in EPC-treated and follicle-co-cultured endometrial organoids. Expression values are row-scaled across samples. **(B)** GSEA comparing enrichment of stage-associated decidualization gene sets between EPC and follicle co-culture. Normalized enrichment scores (NES) are shown; positive NES indicate preferential enrichment in EPC. **(C)** Endest reference-based transcriptional staging of follicle co-culture and EPC samples. Endest scores were used for comparative positioning of the two *in vitro* conditions rather than absolute histological staging. **(D)** Pathway enrichment analysis comparing EPC and follicle co-culture, highlighting decidual maturation programs preferentially associated with EPC and inflammatory/growth-factor and proliferation-associated programs preferentially associated with follicle co-culture. Bars represent NES and shading indicates statistical significance. Statistical significance for the Endest comparison was determined as described in the Materials and Methods.

We next applied the Endest staging model to our transcriptional profiles. Endest was originally developed to capture cyclical transcriptional variation in human endometrial tissue [29]. Given the differences in cellular composition between our *in vitro* samples and whole endometrial biopsies, Endest was used only as a reference-based estimate of the relative transcriptional states of the two models. The coculture model exhibited a relatively earlier transcriptional state than EPC (Figure 5C). Finally, independent pathway enrichment analysis further supported these differences. Decidual maturation programs, including IGFBP1/DKK1-positive cell signatures, cholesterol biosynthesis, IGF receptor binding, and hormone signaling, were preferentially enriched in the EPC group (Figure 5D, Supplemental Table 6). In contrast, inflammatory and growth factor signaling, including cytokine, prostaglandin, EGF, TNFα–NF-κB, and VEGFA-related pathways, as well as proliferation-associated signatures, were enriched in the follicle coculture group. Together, these findings support the notion that the coculture model retains transcriptional features characteristic of an earlier decidualization state.

### Follicle co-culture enables functional evaluation of disrupted decidualization of endometrial organoids

To evaluate the application potential of the follicle–endometrial organoid co-culture model for functional testing of factors that may perturb decidualization, endometrial receptivity, and early pregnancy, we used the PR antagonist RU486 as a proof-of-concept pharmacological challenge. The treatment of RU486 markedly attenuated the morphological changes induced by follicle co-culture, with treated endometrial organoids retaining a more compact morphology compared with organoids co-cultured with luteinized follicles alone (Figure 6A). Consistent with these morphological effects, RU486 significantly reduced the co-culture-induced expression of decidualization markers, including *FOXO1, PRL, DKK1, DCN,* and *REN* (Figure 6B). Together, these results suggest that the follicle–endometrial organoid co-culture system enables functional testing of decidualization, providing a proof of concept for its use as a functional experimental platform to investigate the physiology and pathophysiology of ovarian–endometrial communication and to evaluate the effects of drugs and environmental chemicals on endometrial decidualization, uterine receptivity, and early pregnancy.

**Figure 6.**
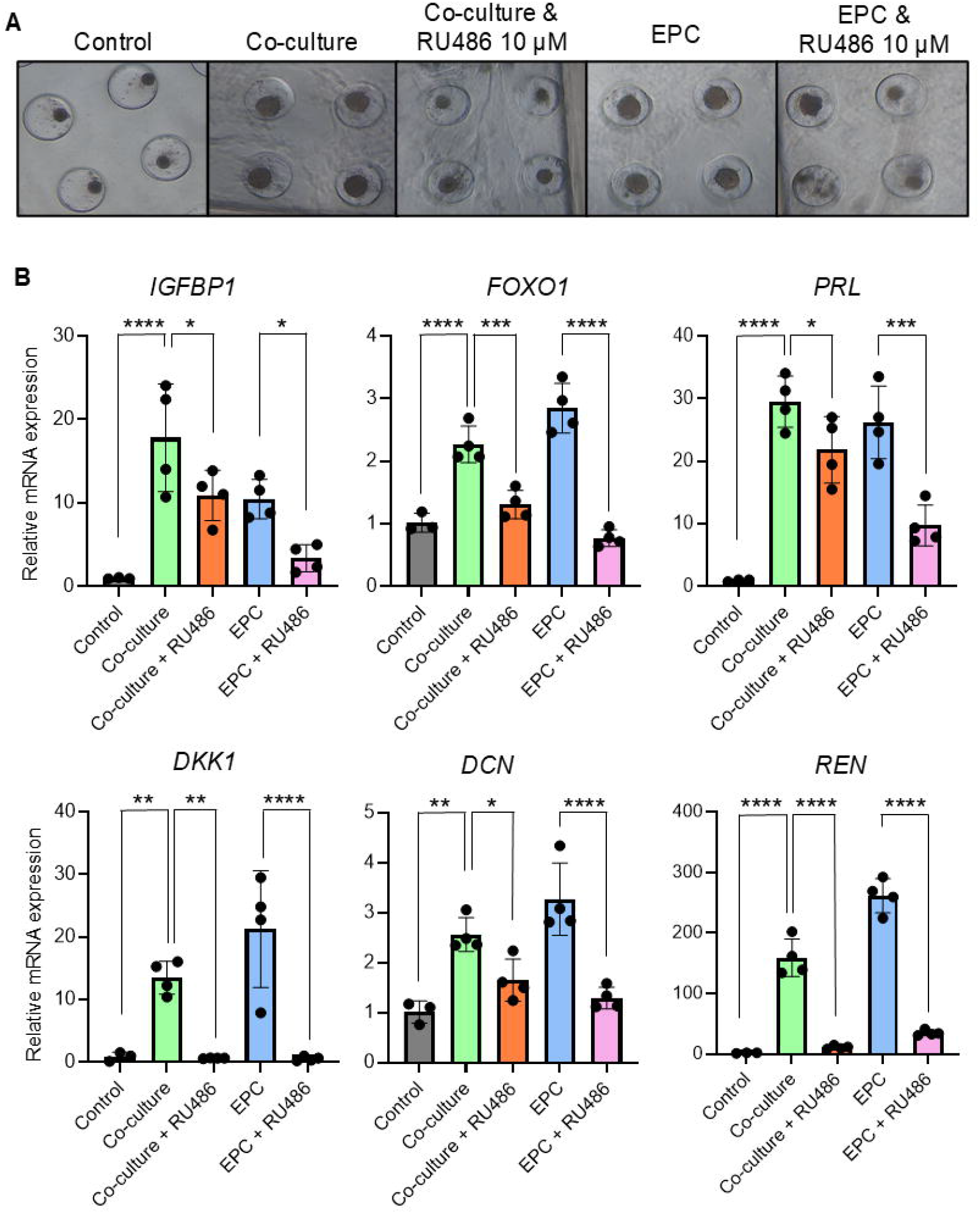
Pharmacological perturbation demonstrates the utility of the follicle–endometrial co-culture model for functional testing of decidualization. Human endometrial organoids were treated with the progesterone receptor antagonist RU486 (10 μM) during the 6-day decidualization period in both follicle co-culture and EPC conditions. **(A)** Representative bright-field images of endometrial organoids from vehicle control, follicle co-culture, follicle co-culture plus RU486, EPC, and EPC plus RU486 groups. **(B)** RT-qPCR analysis of the decidualization-associated genes *IGFBP1, FOXO1, PRL, DKK1, DCN,* and *REN* following treatment. Data are presented as mean ± SD. Statistical significance was determined as described in the Materials and Methods. \**P < 0.05*, \*\**P < 0.01*, \*\*\**P < 0.001*, \*\*\*\**P < 0.0001*.

## Discussion

The present study established a 3D *in vitro* follicle–endometrial organoid co-culture model that enables human endometrial decidualization to be driven by the dynamic secretory activity of the ovarian compartment rather than solely by predefined concentrations of exogenous hormones. Following hCG-induced ovulation and luteinization, the luteinized follicles generated a progressive increase in P4 accompanied by declining E2. Exposure of human endometrial organoids to this changing ovarian environment induced morphological changes and robust expression of decidualization markers. Transcriptomic analyses further demonstrated strong genome-wide concordance between follicle co-culture and conventional EPC-induced decidualization, while revealing distinct transcriptional states between the two conditions. Finally, pharmacological perturbation with RU486 demonstrated that this co-culture model can be used to test functional disruption of endometrial decidualization. Together, these findings provide proof of concept for a multi-organ reproductive NAM that functionally links ovarian activity to human endometrial responses and offers a new experimental framework for investigating ovarian–endometrial communication and evaluating how exogenous exposure to drugs or environmental chemicals perturbs early pregnancy.

A major advance of the present co-culture models is the incorporation of a functional ovarian compartment that generates temporally changing endocrine signals. In the human menstrual cycle, ovarian-derived E2 promotes endometrial proliferation before ovulation. Decidualization is initiated during the postovulatory phase and primarily driven by rising P4 levels and activation of local cAMP signaling [1, 30, 31]. Conventional *in vitro* decidualization models commonly use fixed concentrations of E2, P4, and cAMP to elucidate the molecular mechanisms underlying decidual differentiation [3–6]. However, these approaches do not recapitulate the dynamic secretory changes associated with ovarian function. Importantly, the CL is not only a source of P4 and E2 but also produces steroid metabolites, prostaglandins, relaxin, and multiple vasoactive and angiogenic factors that contribute to implantation, vascular adaptation, placentation, and early pregnancy [2]. Clinical observations further support the importance of a functional CL, as programmed frozen embryo transfer cycles performed in the absence of a CL have been associated with altered maternal cardiovascular adaptation and increased risks of hypertensive disorders of pregnancy[2]. Thus, recapitulating the integrated secretory activity of a functional ovarian compartment may provide biological information that cannot be reproduced by simply adding fixed concentrations of EPC. In our model, the progressive increase in P4 and concurrent decline in E2 following hCG stimulation demonstrate that the ovarian compartment undergoes a dynamic endocrine transition and provides a physiologically motivated context for studying downstream endometrial responses.

The response of human endometrial organoids to co-cultured luteinized follicles was demonstrated across morphological, molecular, and transcriptomic levels. Decidualization involves the transformation of fibroblast-like endometrial stromal cells into enlarged secretory decidual cells, accompanied by extensive transcriptional, metabolic, inflammatory, and tissue-remodeling changes [1, 30]. Consistent with this process, follicle-co-cultured endometrial organoids exhibited increased cellular size and reduced cellular compaction and showed robust induction of canonical decidualization markers, including *IGFBP1, PRL*, and *FOXO1*. Moreover, transcriptomic analysis demonstrated that the response extended well beyond individual markers. Gene-wise changes induced by follicle co-culture and EPC showed strong genome-wide concordance, and a previously defined *in vitro* decidualization gene set was strongly enriched under both conditions. Concordant responses also encompassed multiple biological processes associated with decidualization, including hormone-responsive signaling, immune and inflammatory regulation, lipid and energy metabolism, and cell–cell and cell–matrix interactions. These processes are integral components of decidual transformation and the establishment of a receptive endometrial environment [1, 30, 31]. Thus, the follicle-derived secretory factors reproduce not only selected molecular markers but a broad decidualization-associated transcriptional program.

Despite the broad concordance, follicle co-culture and EPC did not generate identical transcriptional states. EPC-treated organoids preferentially expressed several mid- to late-secretory and first-trimester-associated decidualization markers and showed greater enrichment of pathways associated with decidual maturation, whereas endometrial organoids co-cultured with luteinized follicles retained greater expression of early-secretory-associated genes and inflammatory, growth-factor, vascular, and proliferation-related programs. Reference-based Endest analysis similarly positioned the follicle co-culture group toward a relatively earlier transcriptional state than EPC. These findings are consistent with the dynamic nature of human decidualization, which is not a binary cellular transition but rather involves sequential transcriptional and functional states that support endometrial receptivity, embryo selection, implantation, and subsequent pregnancy [30]. One possibility is that continuous exposure to fixed concentrations of E2, P4, and cAMP drives stromal cells more rapidly toward a mature decidual phenotype, whereas the gradually changing ovarian environment generated by luteinized follicles produces a more progressive decidualization response. However, the differences observed here should not be interpreted simply as evidence that follicle co-culture induces a weaker or incomplete decidual response. Because the luteinized follicular compartment generates a complex and temporally changing mixture of steroids and other secretory products [2], it may also produce a qualitatively different decidualization trajectory from that generated by fixed EPC stimulation. The greater representation of inflammatory, growth-factor, vascular, and proliferative programs in the co-culture condition may therefore reflect both the relative timing of decidualization and the broader signaling environment generated by a functional ovarian compartment. Further time-resolved studies will be required to distinguish these possibilities.

Beyond modeling basic reproductive physiology, an important application of this system is its potential use as a functional NAM to investigate factors that perturb early pregnancy. Advances in 3D human endometrial models have substantially improved the ability to study hormone responsiveness, epithelial–stromal interactions, and decidualization *in vitro* [7, 8]. For example, studies of endometrial epithelial organoids have demonstrated that hormone-regulated epithelial secretions can directly modulate stromal decidualization, highlighting the importance of multicellular and intercompartmental signaling in shaping endometrial responses [8]. Nevertheless, most current endometrial models examine uterine responses independently of a functional ovarian compartment. In the present study, RU486 was therefore used not to establish the well-recognized requirement for PR signaling in decidualization [30, 31], but as a proof-of-concept perturbation to determine whether the integrated model can detect functional disruption of decidualization. RU486 attenuated co-culture-induced decidualization, demonstrating the system’s responsiveness to exogenous perturbation. This capability creates opportunities to investigate both physiological and pathophysiological endometrial decidualization and to evaluate drugs and environmental chemicals that may act directly on the endometrium, alter ovarian function, or disrupt communication between the endometrium and the ovary.

Several limitations of the present proof-of-concept model should be considered. First, because access to functional human ovarian follicles is highly limited, the current system combines murine follicles with human endometrial organoids. The mouse follicular compartment provides a robust and experimentally tractable means to reproduce follicular development, ovulation, luteinization, and dynamic hormone production, but species differences in ovarian hormones and other secretory factors may influence cross-tissue signaling. Future incorporation of human ovarian follicles or stem cell-derived human ovarian models would further improve its translational relevance. Second, the current endometrial organoids predominantly represent the stromal compartment and do not reproduce the full cellular complexity of the human endometrium. Endometrial epithelial cells, immune cells, endothelial cells, and stromal cells interact extensively during early pregnancy[8, 30]. Incorporating these components would therefore enable the model to capture additional dimensions of uterine receptivity and early pregnancy. Finally, future integration with trophoblast or embryo-surrogate models could extend the system beyond decidualization to investigate implantation and early maternal–placental interactions.

In summary, the present study provides proof of concept that a 3D *in vitro* follicle-endometrial organoid co-culture model reproduces ovarian control of endometrial decidualization. This new NAM model provides a foundation for developing human-relevant reproductive models to investigate female reproductive physiology and pathophysiology and to evaluate how therapeutic and environmental exposures perturb uterine receptivity and early pregnancy.

## Author Contributions

Y. Liu, Y. Sun, and T. Zhan contributed to experimental design, data collection and analysis, and manuscript writing; X Liu and J. Zhang contributed to data collection. F. Gao, NC. Douglas, and Q. Zhang contributed to result discussion, data interpretation and manuscript editing; S. Xiao conceived the project, contributed to experimental design, manuscript writing and editing, data analysis and interpretation, and provided final approval of the manuscript.

## Funding and acknowledgments

This work is supported by the National Institutes of Health (NIH) R01ES032144 and OT2OD042764 to S. Xiao and Q. Zhang, R01ES035766 to S. Xiao, Q. Zhang, and NC. Douglas, P30ES005022 and R01HD114750 to S. Xiao.

## Data availability

All associated RNA-seq data, including fastq files and processed raw counts, are available at the Gene Expression Omnibus (GSE325553).

