## Supplemental Table 1 and 2 for "A 3D *in vitro* follicle-endometrial organoid co-culture model recapitulates ovarian control of human endometrial decidualization"

**Supplemental Table 1:** Established decidualization-associated genes and their identified functions in the endometrium

| **Gene** | **Description** | **Ovulatory functions and references** |
| --- | --- | --- |
| *IGFBP1* | Insulin-like Growth Factor-Binding Protein-1 | Decidualization marker and regulation of trophoblast interaction(1, 2) |
| *PRL* | Prolactin | Decidualization marker and local endocrine support(3, 4) |
| *LEFTY* | Left–Right Determination Factor 2 | Negative regulator of decidualization and TGF-β signaling(5) |
| *FOXO1* | Forkhead Box O1 | Master transcriptional regulator of decidualization(6) |
| *DKK1* | Dickkopf WNT Signaling Pathway Inhibitor 1 | Negative modulation of WNT signaling to regulate decidualization and endometrial receptivity(7) |
| *DCN* | Decorin | Regulates extracellular matrix remodeling and growth factor signaling to maintain decidual structure and control trophoblast invasion(8) |
| *REN* | Renin | Local renin–angiotensin signaling to support decidualization and vascular remodeling(9) |
| *SST* | Somatostatin | Local hormonal modulation of decidualization and endometrial receptivity(10) |

**Supplemental Table 2.** Primer sequences of examined genes by RT-qPCR

| **Gene name** | **Sequence** |
| --- | --- |
| *GADPH* | F: CGCTTCGCTCTCTGCTCCTCCTGT  R: GGTGACCAGGCGCCCAATACGA |
| *IGFBP1* | F: AGAGTCGTAGAGAGTTTAGC  R: ACACTGTCTGCTGTGATAA |
| *PRL* | F: CTACATCCATAACCTCTCCTCAG  R: GGGCTTGCTCCTTGTCTTC |
| *LEFTY* | F: GCTGCTCCATGCCGAACACCA  R: CCGCGGAAAGAGGTTCAGCCA |
| *FOXO1* | F: CGAGCTGCCAAGAAGAAA  R: TTCGAGGGCGAAATGTAC |
| *DKK1* | F: GGTATTCCAGAAGAACCACCTTG  R: CTTGGACCAGAAGTGTCTAGCAC |
| *DCN* | F: ATGAAGGCCACTATCATCCTCC  R: GTCGCGGTCATCAGGAACTT |
| *REN* | F: CCGTGATCCTCACCAACTACA  R: ACCCAAACATTGGACGAACCA |
| *SST* | F: ACCCAACCAGACGGAGAATGA  R: GCCGGGTTTGAGTTAGCAGA |

*F: forward; R: reverse
